# Adeno-Associated Virus Mediated Expression of Bcl-xL Attenuates Apoptosis in Fuchs Endothelial Corneal Dystrophy

**DOI:** 10.64898/2026.09.01.748650

**Authors:** Ness Little, Judy Yan, Narisa Dhupar, Stephan Ong Tone

## Abstract

Fuchs Endothelial Corneal Dystrophy (FECD) is characterized by progressive corneal endothelial cell loss and the formation of corneal guttae. Currently, there is a global shortage of donor corneas and new strategies are needed to reduce the need for corneal transplantation. While adeno-associated viruses (AAVs) have the capacity to deliver anti-apoptotic genes to human corneal endothelial cells (CECs), this has not been fully explored as a therapeutic strategy for FECD. In this study, we evaluated the transduction efficiency of self-complementary (sc-) and single-stranded (ss-) AAV2 serotypes in human CECs and ex vivo tissues and assessed whether AAV-mediated expression of Bcl-xL could attenuate apoptosis in FECD. Seventeen scAAV2 serotypes were screened for transduction efficiency in normal human CECs via green fluorescent protein (GFP) expression. The top 4 AAV2 serotypes were further evaluated in FECD cell lines, healthy cadaveric donor specimens, and FECD patient specimens. FECD cell lines were transduced with anti-apoptotic ssAAV2/5-Bcl-xL (AAV2/5-CAG-eGFP-P2A-BCLXL) and treated with etoposide to induce apoptosis. We found that scAAV2/5 demonstrated high transduction efficiency across all normal and FECD cell lines and tissues. We observed that ssAAV2/5-mediated expression of Bcl-xL provided significant protection against etoposide-induced apoptosis in FECD CECs (71.68%±0.69 vs 23.96%±8.88%, *p*=0.018). Our findings show that AAVs have the potential for therapeutic gene delivery to the human corneal endothelium, and that targeting the Bcl-xL mediated apoptotic pathway can be further explored as a therapeutic for FECD.

**Highlights:**

- 17 scAAV2 serotypes were screened in human corneal endothelial cells
- scAAV2/5 can transduce the corneal endothelium with high efficiency
- ssAAV2/5-Bcl-xL offers significant protection against etoposide-induced apoptosis

## 1. Introduction

The human corneal endothelium (CE) is composed of a monolayer of hexagonal corneal endothelial cells (CECs) that are situated on a specialized basement membrane known as Descemet’s membrane (Katikireddy et al., 2016). The CE is the posterior layer of the cornea and plays an essential role in maintaining corneal clarity and normal vision (Benischke et al., 2017). Fuchs Endothelial Corneal Dystrophy (FECD) is an age-related, corneal disorder associated with genetic and environmental factors that primarily affects women and ultimately leads to corneal edema and vision loss (Ong Tone et al., 2021a; Sundin et al., 2006). Currently, FECD is the leading indication of corneal transplantation and is the most common form of corneal dystrophy worldwide, affecting approximately 4% of people over the age of 40 (Afshari, 2006; Chu et al., 2020; Gain et al., 2016; Jurkunas, 2018; Krachmer et al., 1978; Louttit et al., 2012; Minear et al., 2013; Schmedt et al., 2012b; Thompson et al., 2003). FECD pathology includes progressive CEC loss and the formation of excess extracellular matrix (ECM) excrescences, known as guttae (Jun, 2010; Krachmer et al., 1978; Vogt, 1921; Wieben et al., 2012). Ultimately, oxidative stress and mitochondrial dysfunction cause an accumulation of mitochondrial and nuclear DNA damage, leading to cell death (Ong Tone et al., 2021a). While parthanatos and ferroptosis contribute to the cell loss seen in FECD, the intrinsic apoptosis pathway is a major contributor (Saha et al., 2024; Sakakura et al., 2023). Therefore, targeting the intrinsic apoptotic pathway may attenuate cell death in FECD.

Bcl-2 family proteins play an important role in the regulation of the intrinsic apoptosis pathway and are therefore interesting targets for the therapeutic prevention of apoptosis (Jan and Chaudhry, 2019). Bcl-xL, or B-cell lymphoma extra large, is a Bcl-2 family anti-apoptotic protein that plays a critical role in the prevention of apoptosis. It works to inhibit the loss of mitochondrial outer membrane potential (MOMP) and cytochrome C from the intermembrane space and can protect CECs from apoptosis (Barcia et al., 2007; Jan and Chaudhry, 2019; Wang et al., 2006). The anti-apoptotic effects of Bcl-xL makes it a compelling target for anti-apoptotic gene therapy, particularly in the context of the apoptotic dysregulation seen in FECD.

Bcl-xL has been investigated as a neuroprotective factor in a number of neurodegenerative conditions, which share striking cellular and molecular similarities to FECD (Garrity-Moses et al., 2005; Zhu et al., 2014). Bcl-xL overexpression has been shown to significantly decrease apoptotic cell death in in vitro and in vivo rat models of amyotrophic lateral sclerosis (ALS), Alzheimer’s, and Parkinson’s diseases (Garrity-Moses et al., 2005; Kaspar et al., 2002; Matsuoka et al., 1999). Overexpression of Bcl-xL has also been found to preserve mitochondrial structure and function, aid in the regulation of oxidative stress and intrinsic apoptosis, and increase mitochondrial resistance to apoptosis (Mas-Bargues et al., 2025). Together, these studies support the use of Bcl-xL gene transfer as a therapeutic strategy for degenerative diseases and FECD.

Despite a significant need to develop improved interventions to halt or reverse FECD pathogenesis, limited studies have explored using adeno-associated viruses (AAVs) to deliver anti-apoptotic genes to the corneal endothelium, and they have not been fully explored as a therapeutic strategy for FECD (Barcia et al., 2007). AAVs are a type of DNA viral vector that can transduce both dividing and non-dividing cells and are currently being used in human clinical trial and gene therapies (Bastola et al., 2020; Daya and Berns, 2008; George et al., 2000; Joyce, 2003; Kuot et al., 2017). AAVs have many advantages that confer a robust safety profile, including that they elicit a mild immune response, are episomal in nature, and avoid random integration into the host genome (Bastola et al., 2020). Additionally, AAVs are non-pathogenic viral vectors and drive long term gene expression (Mohan et al., 2011). While over 100 AAV serotypes are known to exist, twelve (AAV serotypes 1-12) have been identified as major human serotypes, and nine (AAV serotypes 1-9) have been extensively studied for gene therapy uses (Cui et al., 2022; Mohan et al., 2011). AAV2 has been used widely in many clinical and preclinical trials in the eye and is the first FDA-approved gene therapy to treat a genetic retinal disorder (Leber congenital amaurosis) (Keeler and Flotte, 2019). In addition to AAV2’s broad tropism and widespread use, AAV2 has been shown to transduce human CECs with high efficiency, making it a promising serotype for the delivery of gene therapies to the CE (Gruenert et al., 2016).

In this study, we screened 17 self complementary (sc-) AAV2 serotypes for their ability to transduce human CECs (HCEC-21T) and investigated 4 serotypes with high transduction efficiency in an additional human CEC line (HCEC-SVN2-67F) and healthy cadaveric donor cornea tissues. We subsequently investigated the transduction efficiency of 2 AAV serotypes (scAAV2/2 and scAAV2/5) in FECD CECs, and FECD patient specimens. We then investigated the ability of single-stranded (ss-) AAV2/5-mediated expression of the anti-apoptotic gene Bcl-xL to attenuate etoposide-induced apoptosis in FECD.

## 2. Materials and Methods

### 2.1. Human Tissues and Ethical Statements

Informed consent from patients undergoing surgery for FECD was obtained prior to tissue collection. Surgically removed and healthy cadaveric donor tissues were stored in Optisol-GS (Bausch & Lomb) at 4°C for 0–18 days prior to use. Normal tissues from cadaveric donors were provided by The Eye Bank of Canada or Eversight (Chicago, Illinois). See Table 1 for cadaveric donor and patient characteristics. This study was carried out according to the tenets of The Declaration of Helsinki and was approved by the Sunnybrook Health Sciences Centre Research Ethics Board (REB#5070 and REB#5187).

**Table 1.** Patient and Donor Characteristics.

| Ocular Health | Experimental Usage | Age | Sex | Cause of Death | Lens Type | Additional Co-morbidities | Time to Preservation (hours:min) | Preservation to Processing (days) |
| --- | --- | --- | --- | --- | --- | --- | --- | --- |
| Normal | Bulk RNA-seq | 74 | M | Other, Respiratory Arrest | Phakic | COPD, Pneumonia | 8:30 | 16 |
| Normal | Bulk RNA-seq | 71 | F | Anoxia | Unknown | PMH, Diabetes, Nephrolithiasis, Osteomyelitis, Vascular Dementia, CVA, CHF | 22:20 | 15 |
| Normal | Bulk RNA-seq | 76 | M | Other | Unknown | HTN, Dyslipidemia, Diabetes, Alcoholism, Depression, Chronic Cord Dysfunction, Anoxic Brain Injury | 17:45 | 6 |
| FECD | Bulk RNA-seq | 75 | M | Other, CVA | Pseudophakic | CAD, Cancer, Arthritis, Diabetes, Dyslipidemia, | 8:18 | 4 |
| FECD | Bulk RNA-seq | 69 | F | Other | Unknown | Spinal Stenosis, Macular Degeneration | 15:03 | 5 |
| FECD | Bulk RNA-seq | 66 | M | Myocardial Infarction | Phakic | Acute Hypoxemic Respiratory Failure, Cardiac Arrest, Cardiac Disease, Dysphagia, Schizophrenia, Seizures, Tetraplegia/Quadriplegia, TIA | 2:27 | 14 |
| Normal | scAAV2/2 | Unknown | Unknown | Unknown | Unknown | Unknown | Unknown | 18 |
| Normal | scAAV2/2 | 55 | M | CVA/Stroke | Unknown | Unknown | 10:00 | 0 |
| Normal | scAAV2/2 | 51 | F | Anoxia | Phakic | Anoxia, Acute Bacterial Sinusitis, Depression, Anxiety, Seizures, Endocervical Curettage, Arthritis, Pyelonephritis | 7:40 | 11 |
| Normal | scAAV2/5 | 55 | M | CVA/Stroke | Unknown | Unknown | 10:00 | 0 |
| Normal | scAAV2/5 | 9 | F | Anoxia | Phakic | Asthma, Allergies | 3:42 | 18 |
| Normal | scAAV2/5 | 32 | M | Other, OD | Unknown | OD, Asthma, Seizure, Depression | 16:40 | 4 |
| Normal | scAAV2/6 | 46 | F | Other, Unknown | Phakic | Diabetes, HTN, Hyperlipidemia | 23:58 | 6 |
| Normal | scAAV2/6 | 9 | F | Anoxia | Phakic | Asthma, Allergies | 3:42 | 18 |
| Normal | scAAV2/6 | 32 | M | Other, OD | Unknown | OD, Asthma, Seizure, Depression | 16:40 | 4 |
| Normal | scAAV2/DJ | 46 | F | Other, Unknown | Phakic | Diabetes, HTN, Hyperlipidemia | 23:58 | 6 |
| Normal | scAAV2/DJ | 74 | F | Other, Anoxia | Unknown | PMH, COPD, T2DM, Fibromyalgia, HTN, Chronic Kidney Disease, Bilateral Renal Artery Stenosis with Atrophic Left Kidney, Gout, CHF | 9:18 | 4 |
| Normal | scAAV2/DJ | 29 | M | Other, OD | Phakic | Acute Hypoxemic Respiratory Failure, AKI, Anoxic Brain Injury, Cardiac Arrest, Cerebral Edema, Metabolic Acidosis, Seizures, Ischemic Hepatitis | 2:36 | 11 |
| FECD | scAAV2/2 | 74 | M | Other, GI Bleed | Phakic | GI Bleed, HCC, Lung Cancer, T2DM, Dyslipidemia, HTN | 22:57 | 4 |
| FECD | scAAV2/2 | 89 | F | n/a | Pseudophakic | HTN, Rheumatoid Arthritis, Asthma, Bronchitis | 0 | 0 |
| FECD | scAAV2/2 | 75 | F | n/a | Pseudophakic | HTN, Hypothyroid | 0 | 0 |
| FECD | scAAV2/5 | 74 | M | Other, GI Bleed | Phakic | GI Bleed, HCC, Lung Cancer, T2DM, Dyslipidemia, HTN | 22:57 | 4 |
| FECD | scAAV2/5 | 73 | M | n/a | Phakic | Hyperlipidemia, Stroke | 0 | 1 |
| FECD | scAAV2/5 | 65 | F | n/a | Phakic | Rheumatoid Arthritis, Psoriasis | 0 | 1 |

AKI = Acute kidney injury; CAD = Coronary artery disease; CHF = Congestive heart failure; COPD = Chronic obstructive pulmonary disease; CVA = Cerebrovascular accident; GI = Gastrointestinal; HCC = Hepatocellular carcinoma; HTN = Hypertension; OD = Overdose; PMH = Progressive macular hypomelanosis; T2DM = Type 2 diabetes mellitus; TIA = Transient ischemic attack
All tissues preserved in Optisol-GS prior to processing

### 2.2. AAV Production

Packaging of AAV vectors was done at the CNP Viral Vector Core (Canadian Neurophotonics Platform Viral Vector Core Facility (RRID:SCR_016477)). Viral constructs for transfection studies were packaged in self-complementary AAV (scAAV) capsids, whereas constructs for anti-apoptosis studies were packaged in single-stranded AAV (ssAAV) capsids.

During the packaging process for the ssAAV2/5 constructs, Bcl-xL (from mCerulean3-BclXL-pEGFP-C1; Addgene #177408) was cloned into the ssAAV with eGFP. To ensure comparable levels of expression, a P2A peptide sequence was inserted between Bcl-xL and eGFP (“ssAAV2/5-Bcl-xL” (AAV2/5-CAG-eGFP-P2A-BCLXL) and “ssAAV2/5” (AAV2/5-CAG-eGFP); Canadian Neurophotonics Platform Viral Vector Core Facility).

### 2.3. Cell Culture

Immortalized normal and FECD human corneal endothelial cell (CEC) lines were graciously provided by Dr. Ula Jurkunas at The Schepens Eye Research Institute in Boston, Massachusetts (Ashraf et al., 2025; Ong Tone et al., 2021b; Schmedt et al., 2012a). CECs were isolated from a 54-year-old female (FECD-SV-54F-73) and a 61-year-old male (FECD-SVF6-61M) undergoing endothelial keratoplasty for FECD, as well as a 21-year-old male (HCEC-21T) (Schmedt et al., 2012a) and a 67-year-old female (HCEC-SVN2-67F) cadaveric donor (Ashraf et al., 2025).

Cells were cultured in Chen’s Media consisting of Opti-MEM media (ThermoFisher Scientific, Waltham, MA) supplemented with 200 mg/L CaCl_2_ (Millipore Sigma, Oakville, Ontario), 0.08% chondroitin sulfate (Millipore Sigma, Oakville, Ontario), 50 μg/mL gentamicin (ThermoFisher Scientific, Waltham, MA), 1X antibiotic/antimycotic (Wisent, St. Bruno, QC), 66 μg/mL bovine pituitary extract (Gemini, West Sacramento, CA), 5 ng/mL epidermal growth factor (Millipore Sigma, Oakville, Ontario) and 8% fetal bovine serum (ThermoFisher Scientific, Waltham, MA). Cell culture plates were coated with fibronectin coating mix (AthenaES, Baltimore, MD) prior to cell seeding and incubated at 37℃ and 5% CO_2_.

### 2.4. Bulk RNA Sequencing in Normal and FECD Donor CE

Descemet’s membrane and the cornea endothelium were collected from normal (n=3) and FECD cadaveric donors (n=3). Total RNA was isolated from each tissue using the PureLink RNA Micro Kit (ThermoFisher Scientific, Waltham, MA) according to the manufacturer’s protocol. Samples were sent to the Genomics Core Facility at Princess Margaret Cancer Centre in Toronto, Canada and the quantity of total RNA were determined using an Agilent 2100 Bioanalyzer with an RNA 6000 Pico Kit (Agilent Technologies, Santa Clara, CA). The quality of total RNA was assessed by RNA integrity number (RIN) using the Agilent 2100 Expert Software (Agilent Technologies). The RNA-Seq libraries for next-generation sequencing (NGS) were generated with Illumina Stranded Total RNA Ribo Zero Plus kit according to the manufacturer’s instructions. Sequencing was carried out with the NovaSeq 6000 using a 100-cycle paired read protocol and multiplexing to obtain ∼40 million reads/sample.

The read quality on the raw sequencing data was checked using FASTQC v.0.11.5. (Omar et al., 2020). The raw sequencing data were aligned to the human genome (GRCh38, Homo_sapiens.GRCh38.84.gtf) using the HISAT2 v2.2.1. (Kim et al., 2019). The accessory software tool for the alignment stage includes Samtools v1.17. (Li et al., 2009). Transcript assembly was completed using StringTie v2.1.4. (Pertea et al., 2015). HTSeq (v0.11.0) toolkit was utilized to access the htseq-count tool which pre-processed the RNA-seq alignments and generated read counts. Alignment quality control was completed using Picard v2.10.9 (“Picard Toolkit”) and MultiQC v1.7. (Ewels et al., 2016). Genes of interest were identified using a filter (FPKM > 5).

### 2.5. scAAV2 Screening in Cell Lines

Seventeen different self-complementary AAV2 serotypes (scAAV2/1, 2/2, 2/4, 2/5, 2/6, 2/7, 2/8, 2/9, 2/Rh10, 2/DJ, 2/DJ8, 2/Retro, 2/php.B, 2/php.eB, 2/php.S, 2/php.N, and 2/php.V1) expressing GFP under a CAG promoter (AAV-scCAG-eGFP; Canadian Neurophotonics Platform Viral Vector Core Facility) were screened in normal HCEC-21T cells to maximize transduction and minimize vector toxicity.

Briefly, HCEC-21T cells were seeded in 24-well plate at a density of 50,000 cells/well and incubated over night at 37°C. Chambers were washed with 1X PBS, transduced with 1x10^10^ vector genomes of AAV (MOI of 200,000 assuming 50,000 cells/chamber), and incubated overnight at 37°C. Virus-media was removed the following day and replaced with fresh Chen’s media. Cells were incubated for an additional 48 hours at 37°C. Cells were washed with 1X PBS, fixed with 4% paraformaldehyde (ThermoFisher Scientific, Waltman, MA) for 20 min at room temperature and permeabilized with 0.2% Triton-X (Biorad, Mississauga, Ontario). Cells were incubated with 100μg/mL Ribonuclease A (Sigma Aldrich, Saint Louis, MO) for 1 hour at 37°C. Nuclei were stained with propidium iodide (1:3000, Sigma Aldrich, Saint Louis, MO) for five minutes at room temperature in a darkened chamber. Images were obtained throughout using the IncuCyte S3 Live Cell Analysis System (Sartorius). Transduction was quantified using ImageJ as the percentage of total cells expressing GFP from a series of 4-6 representative images per condition. Selected scAAV2 serotypes (scAAV2/2, 2/5, 2/6, 2/DJ) were validated in HCEC-SVN2-67F, FECD-SV-54F-73, and FECD-SVF6-61M cell lines following the same methodology.

### 2.6. Ex vivo Tissue Transduction

The top four performing scAAV2 serotypes (scAAV2/2, 2/5, 2/6, 2/DJ) were first tested in normal donor cornea endothelium. Corneal endothelium and Descemet’s membrane (DM) was isolated from cadaveric donor corneas. Briefly, the corneal endothelium-DM complex was peripherally dissected from the underlying stroma and peeled away using tying forceps and place in Chen’s media. Ex vivo tissues were transduced with 1x10^10^ vector genomes/tissue and incubated in Chen’s media at 37 °C. After 96 hours, media was replaced with fresh Chen’s media. Tissues were incubated for an additional 8-10 days at 37°C. Specimens were fixed with 4% paraformaldehyde for 20 minutes and permeabilized using 0.2% Triton-X for ten minutes at room temperature. Tissues were gently unfolded using 1% hyaluronic acid onto a glass slide and counterstained with VECTASHEILD mounting media with DAPI (Vector Laboratories, Newark, CA). Images were taken with a Leica S DMi8 inverted fluorescence microscope at 20x magnification. Transduction was quantified with ImageJ as the percentage of total cells expressing GFP. A total of 10-15 images were quantified per tissue. FECD ex vivo surgical specimens from patients undergoing endothelial keratoplasty were transduced with scAAV2/2 and scAAV2/5 following the same methodology.

### 2.7. scAAV2 Transduction and Apoptosis Assay

scAAV2/2 and scAAV2/5 were further investigated using HCEC-21T, HCEC-SNV2-67F, and FECD-SV-54F-73 cell lines to determine if scAAV2 infection increases in vitro apoptosis. Briefly, cells were seeded in 24-well plate at a density of 50,000 cells/well and incubated overnight at 37°C. Cells were washed with 1X PBS and transduced with each scAAV2 serotype (1x10^10^ vector genomes) in Chen’s media containing 0.5µM IncuCyte Caspase-3/7 Red Dye (Sartorius), along with controls. Twenty-four hours post-AAV addition, virus and dye were removed and replaced with fresh Chen’s media containing 0.5µM IncuCyte Caspase-3/7 Red Dye. Cells were imaged at 20x magnification every two hours throughout the experiment using the IncuCyte. Apoptosis was quantified 24 hours post AAV addition using ImageJ as the percentage of the total cells expressing caspase-3/7 from 4-5 representative images per well.

### 2.8. ssAAV2/5 Transduction and Apoptosis Assay

FECD-SV-54F-73 cells were plated in a 24-well plate at a density of 50,000 cells and incubated overnight at 37°C. Cells were washed with 1X PBS and transduced with 1x10^10^ vector genomes ssAAV2/5-Bcl-xL, ssAAV2/5, or No AAV in Chen’s media containing 0.5µM IncuCyte Caspase-3/7 Red Dye. Twenty-four hours post-AAV addition, virus and dye were removed and replaced with fresh Chen’s media containing 0.5µM IncuCyte Caspase-3/7 Red Dye and either 17µM etoposide or 1X PBS. Cells were imaged throughout the experiment using the IncuCyte. Apoptosis was quantified 24 hours post-etoposide addition in ImageJ as the percentage of total cells expressing caspase-3/7 dye from 4-5 representative images per well.

### 2.9. Statistical Analysis

Statistical analysis was conducted using R (R Core Team, 2021) and the Tidyverse package (Wickham, 2016). The normality of the distribution of the results was determined using the Shapiro-Wilk test and Q-Q plots. Statistical significance was determined using Welch’s 2-sample t-test, the Wilcoxon Rank Sum Test, or either a one-way or two-way analysis of variance (ANOVA) followed by a Tukey’s HSD post-hoc test. A p-value of <0.05 was considered statistically significant.

## 3. Results

### 3.1. Bulk RNA Sequencing in Normal and FECD Donor CE

Fourteen genes have been identified by multiple independent groups as being associated with AAV entry for various serotypes (*ACP2, ATP2C1, ATP6V0A2, B3GAT3, COG7, C16orf62, EXT1, EXT2, GPR108, KIAA0319L, NDST1, RGP1, RNF121*, and *TM9SF2*) (Meyer and Chapman, 2022). To identify whether these genes are expressed in the CE, we performed bulk RNA sequencing on human corneal endothelium isolated from cadaveric corneas without known corneal pathology (N=3) and cadaveric corneas with FECD (N=3). Of these 14 genes, 12 were expressed in our dataset (*ACP2, ATP2C1, B3GAT3, C16orf62, EXT1, EXT2, GPR108, KIAA0319L, NDST1, RGP1, RNF121*, and *TM9SF2*) (Figure 1) (Meyer and Chapman, 2022). No significant differences were seen in FPKM (fragments per kilobase of transcript per million fragments mapped) levels of each of these 12 genes between normal and FECD donor tissues (*ACP2* (38.74±10.40 and 34.10±7.11, respectively), *ATP2C1* (18.37±4.68 and 19.44±5.73), *B3GAT3* (37.97±17.20 and 35.13±2.34), *C16orf62* (27.69±1.71 and 26.93±6.62), *EXT1* (8.43±0.81 an 14.30±6.58), *EXT2* (41.83±6.39 and 47.55±9.81), *GPR108* (45.48±2.45 and 36.51±8.36), *KIAA0319L* (29.14±1.11 and 26.87±4.10), *NDST1* (12.72±3.68 and 19.86±9.76), *RGP1* (49.74±11.69 and 36.70±7.18), *RNF121* (8.86±0.38 and 8.71±2.45), *TM9SF2* (67.37±6.59 and 73.41±13.74); *p*=0.425).

**Figure 1.**
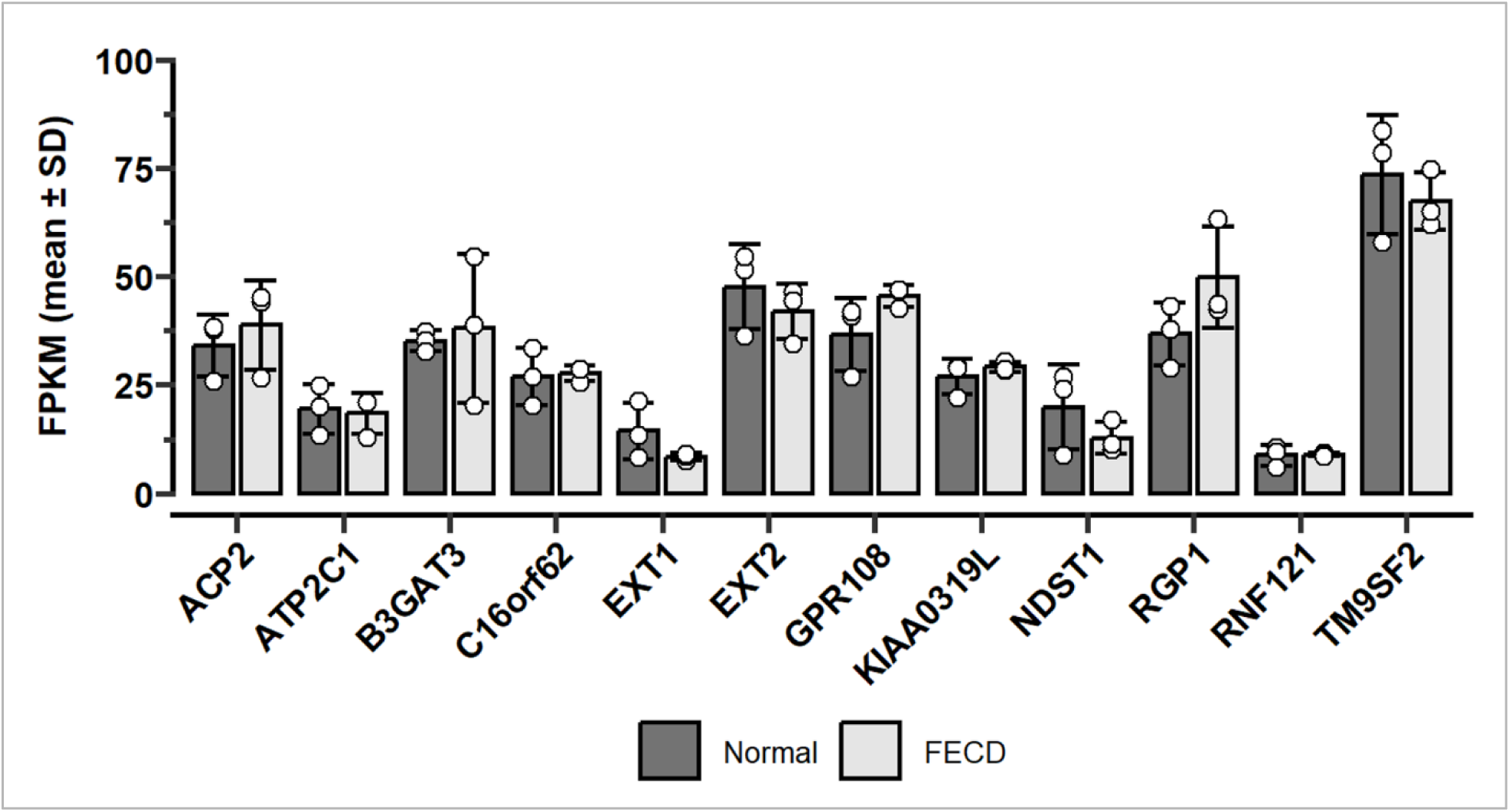
Gene expression levels of AAV binding genes in the human corneal endothelium from healthy and Fuchs endothelial corneal dystrophy specimens. Twelve AAV binding genes were identified in the bulk RNA-seq dataset of normal donor and FECD CE specimens (*ACP2, ATP2C1, B3GAT3, C16orf62, EXT1, EXT2, GPR108, KIAA0319L, NDST1, RGP1, RNF121*, *TM9SF2*). No significant differences were seen in the FPKM levels (*p*=0.425). Statistical analysis was conducted using 2-way ANOVA. AAV = adeno-associated virus, FPKM = fragments per kilobase of transcript per million fragments mapped.

### 3.2. scAAV2 Screening in Human CECs

Given that AAV2 has been used in clinical and preclinical trials in the eye, the evidence that it can transduce human CECs, and the expression of *KIAA0319L* (also known as AAV receptor - AAVR, and critical for AAV2 transduction) in human CECs, we explored the ability of different AAV2 serotypes to transduce human CECs (Gruenert et al., 2016; Keeler and Flotte, 2019; Pillay et al., 2016). Seventeen scAAV2 serotypes expressing GFP under a CAG promoter (AAV-scCAG-eGFP) were screened in normal human CEC line (HCEC-21T) for transduction efficiency via GFP expression, and we identified the top four scAAV2 serotypes with the highest transduction (scAAV2/2 (63.81%±4.65), scAAV2/5 (37.95%±14.17), scAAV2/6 (33.61%±14.13), and scAAV2/DJ (37.64%±11.77)) (Figure 2A, Supplemental Figure 1, Supplemental Figure 2A).

**Figure 2.**
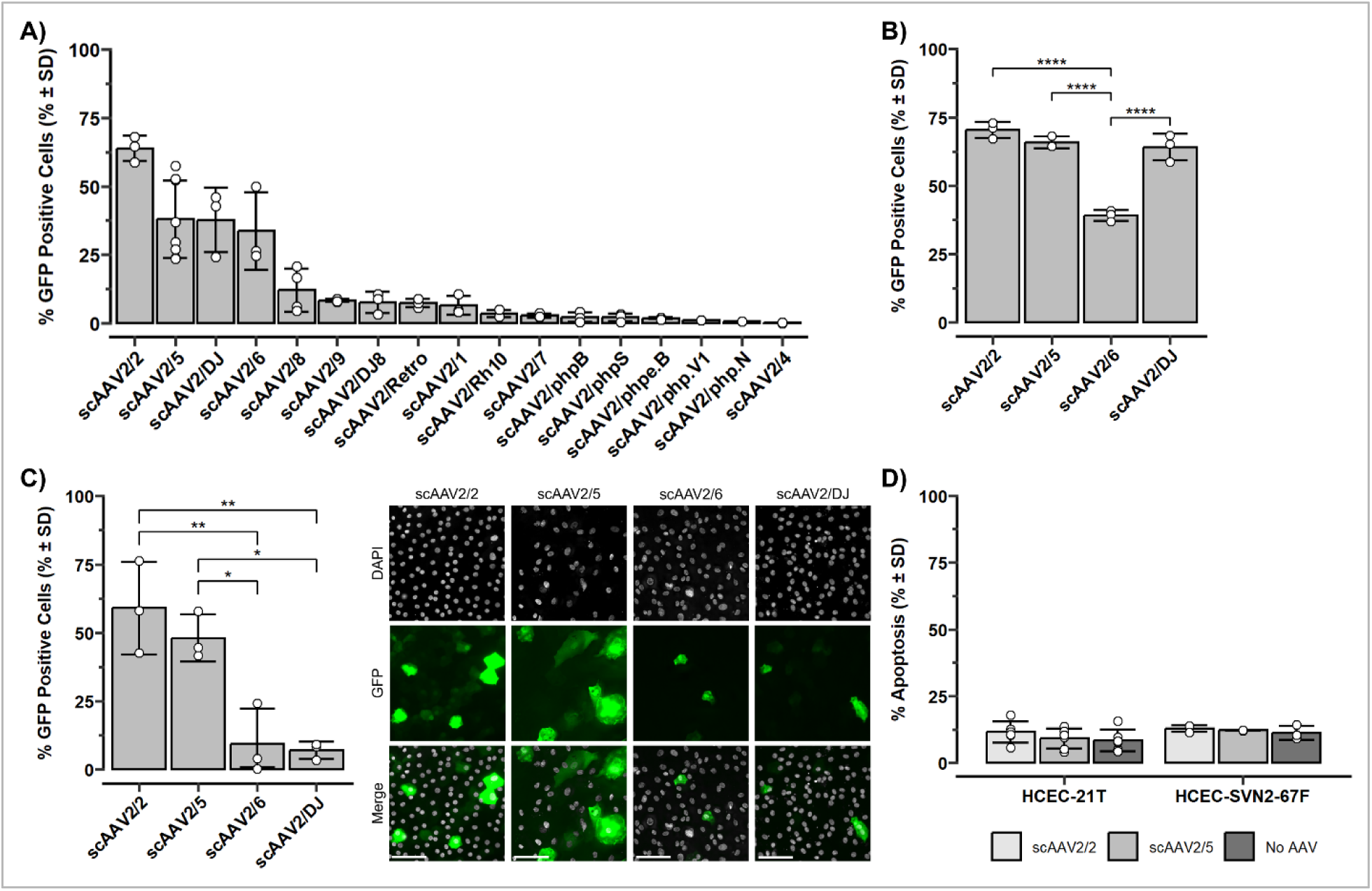
scAAV2/2 and scAAV2/5 transduce human corneal endothelial cell lines and normal cadaveric donor ex vivo corneal tissues. **(A)** Seventeen scAAV2 serotypes were screened in a human corneal endothelial cell line (HCEC-21T) for transduction efficiency and quantified as the percentage of total cells expressing GFP. **(B)** scAAV2/2, scAAV2/5, scAAV2/6, and scAAV2/DJ transduction in another human corneal endothelial cell line (HCEC-SVN2-67F). ANOVA with Tukey’s HSD post-hoc test was performed. **(C)** Left: Quantification of scAAV2/2, scAAV2/5, scAAV2/6, and scAAV2/DJ transduction in normal cadaveric donor tissues. Transduction efficiency was quantified as the percentage of total cells expressing GFP. Statistical analysis was performed using ANOVA with Tukey’s HSD. Right: Representative images of scAAV2/2, scAAV2/5, scAAV2/6, and scAAV2/DJ in normal tissues. Nuclei were stained with DAPI. Scale bar = 100 μm. **(D)** Quantification of apoptosis in HCEC-21T and HCEC-SVN2-67F cells transduced with scAAV2/2 and scAAV2/5. No significant differences were seen in the levels of apoptosis between the transduced and non transduced cells (*p*=0.863). Apoptosis was quantified as the percentage of total cells expressing Caspase-3/7. Statistical analysis was performed using 2-way ANOVA. All experiments were done using N=3. AAV = adeno-associated virus, sc = self-complimentary, GFP = green fluorescent protein, PI = propidium iodide. \**p* ≤ 0.05, \*\**p* ≤ 0.01, \*\*\**p* ≤ 0.001.

### 3.3 scAAV2 Transduction in Human CECs

To further characterize transduction efficiencies between scAAV2/2, 2/5, 2/6, and 2/DJ we transduced an additional human CEC line derived from a healthy cadaveric donor (HCEC-SVN2-67F). We observed high transduction with scAAV2/2 (70.39%±2.98), scAAV2/5 (65.78%±2.17), and scAAV2/DJ (64.10%±4.89) in HCEC-SVN2-67F cells (Figure 2B, Supplemental Figure 2B). scAAV2/6 showed significantly lower transduction (39.05%±1.95) compared to the other four serotypes (*p*_scAAV2/2-scAAV2/6_=0.00001, *p*_scAAV2/5-scAAV2/6_=0.00003, *p*_scAAV2/2-scAAV2/DJ_=0.00005, respectively).

### 3.4. scAAV2 Transduction in Normal Ex Vivo Tissues

We next investigated transduction efficiency of these top 4 scAAV2 serotypes (scAAV2/2, 2/5, 2/6, 2/DJ) in ex-vivo corneal endothelium isolated from healthy cadaveric donor corneas and found that scAAV2/2 (59.03%±16.90) and scAAV2/5 (48.03%±8.61) resulted in the highest transduction efficiency (Figure 2C). Both scAAV2/2 (*p*_scAAV2/2-scAAV2/6_=0.003, *p*_scAAV2/2-scAAV2/DJ_=0.003) and scAAV2/5 (*p*_scAAV2/5-scAAV2/6_=0.015, *p*_scAAV2/5-scAAV2/DJ_=0.011) yielded a significantly higher percentage of GFP positive cells compared to scAAV2/6 (9.44%±12.87) and scAAV2/DJ (7.01%±3.08) (Figure 2C).

### 3.5. scAAV2 Transduction is Not Associated with Increased Apoptosis In Vitro

We next investigated whether scAAV2/2 and scAAV2/5 transduction resulted in increased apoptosis in normal human CECs. No significant differences in apoptosis were observed with scAAV2/2 or scAAV2/5 transduction compared to baseline in HCEC-21T (No AAV=8.43%±4.02, scAAV2/2=11.70%±3.96, scAAV2/5=9.14%±3.77) or HCEC-SVN2-67F (No AAV=11.27%±2.60, scAAV2/2=12.82%±1.27, scAAV2/5=12.13%±0.10) cell lines (*p*=0.863) (Figure 2D, Supplemental Figure 3).

### 3.6. scAAV2/2 and scAAV2/5 Transduction in FECD CECs

We next investigated the ability of scAAV2/2 and scAAV2/5 to transduce two FECD CEC lines derived from FECD patients (FECD-SV-54F-73 and FECD-SVF6-61M). We observed good transduction efficiencies of both scAAV2 serotypes in both cell lines, and no significant differences in transduction efficiencies were observed between scAAV2/2 and scAAV2/5 in FECD-SV-54F-73 (44.90%±20.66 and 33.22%±13.50, respectively; *p*=0.466) or FECD-SVF6-61M cells (54.49%±5.19 and 41.97%±10.27, respectively; *p*=0.157) (Figure 3A, Supplemental Figure 2C,D). We next investigated whether scAAV2/2 and scAAV2/5 transduction resulted in increased apoptosis in FECD. No significant differences in apoptosis were observed with scAAV2/2 or scAAV2/5 transduction compared to baseline in FECD-SV-54F-73 (No AAV=15.36%±11.82, scAAV2/2=9.12%±3.28, scAAV2/5=11.85%±8.16) (*p*=0.683) (Figure 3B, Supplemental Figure 3).

**Figure 3.**
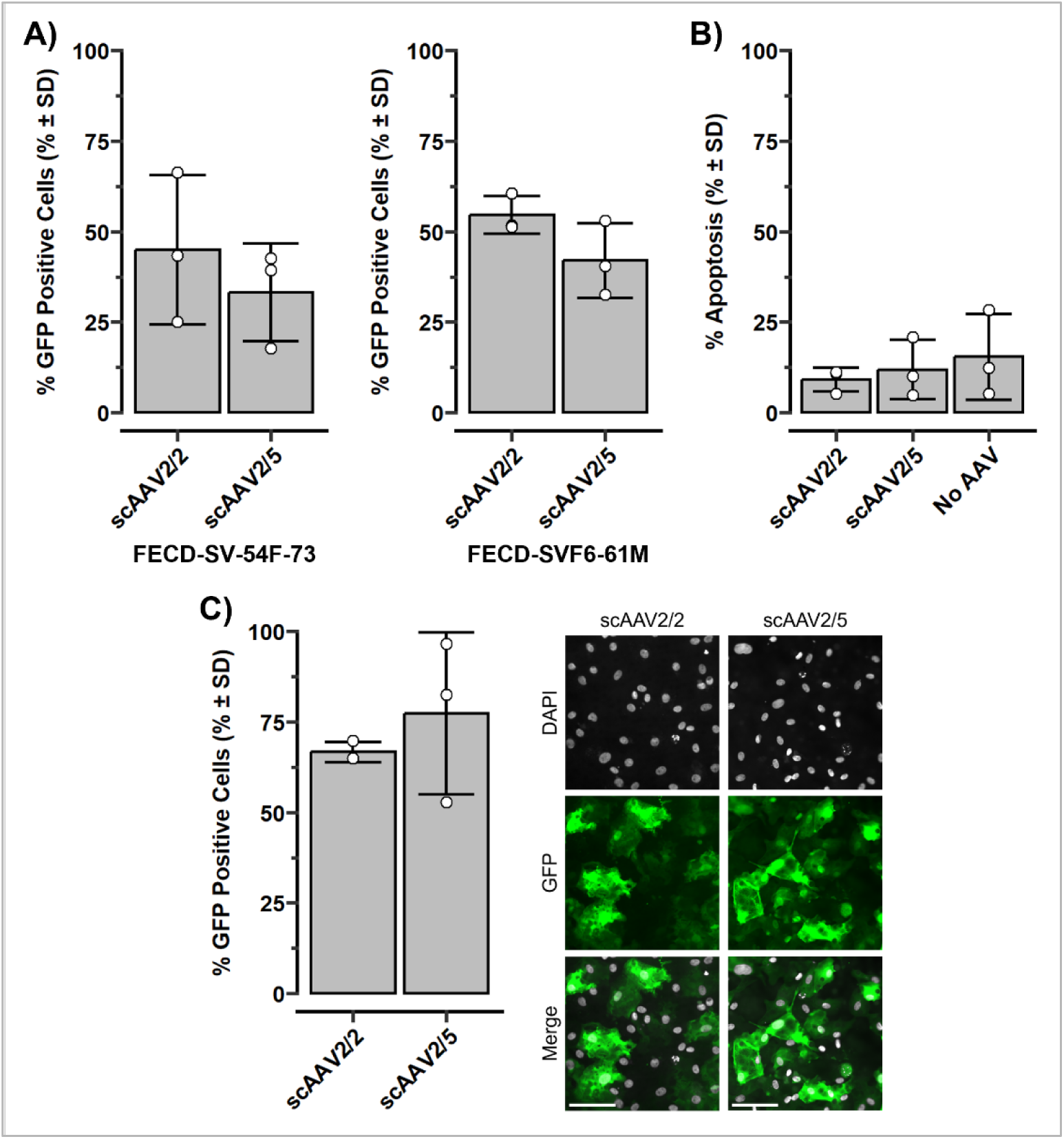
scAAV2/2 and 2/5 show high transduction efficiency in FECD. **(A**) scAAV2/2 and scAAV2/5 transduction in FECD-SV-54F-73 (Left) and FECD-SVF6-61M (Right) cells. Transduction efficiency was quantified as the percentage of total cells expressing GFP. No significant differences were seen in between the serotypes in either cell line (*p*_FECD-SV-54F-73_=0.466, *p*_FECD-SVF6-61M_=0.157). Statistical analysis using Welch’s 2-sample t-test. **(B)** Quantification of apoptosis in FECD-SV-54F-73 HCECs. No significant differences were seen in the levels of apoptosis between the transduced and non transduced cells (*p*=0.683). Apoptosis was quantified as the percentage of total cells expressing Caspase-3/7. Statistical analysis was performed using one-way ANOVA. **(C)** Left: Quantification of scAAV2/2 and scAAV2/5 transduction in FECD ex vivo surgical specimens. No significant differences were seen between the serotypes (*p*=0.700). Statistical analysis was performed using Wilcoxon Rank Sum Test. Right: Representative images showing the GFP expression in FECD ex vivo surgical specimens transduced with scAAV2/2 and scAAV2/5. Nuclei were stained using DAPI. Scale bar = 100 μm. All experiments were done using N=3. AAV = adeno-associated virus, sc = self-complimentary, GFP = green fluorescent protein.

### 3.7. scAAV2/2 and scAAV2/5 Transduction in FECD Ex Vivo Tissues

We next investigated the transduction efficiencies of scAAV2/2 and scAAV2/5 in ex-vivo corneal endothelium isolated from FECD patients undergoing endothelial keratoplasty. We found that both scAAV2/2 (66.60%±2.75) and scAAV2/5 (77.25%±22.34) demonstrated high transduction efficiency and with no significant differences between the serotypes (*p*=0.700) (Figure 3C).

### 3.8. ssAAV2/5-Bcl-xL attenuates etoposide-mediated apoptosis in FECD CECs

To investigate the ability of AAV2 as a delivery vector for gene therapy for FECD, we designed a strategy to introduce the anti-apoptotic gene Bcl-xL to FECD CECs and test its ability to protect from apoptosis. We induced apoptosis with etoposide, a topoisomerase II inhibitor previously used to activate the intrinsic apoptosis pathway in human CECs (Barcia et al., 2007; Fuchsluger et al., 2011a; Staehlke et al., 2024). Furthermore, apoptosis is a key downstream pathway that is activated in FECD and leads to cell death (Jurkunas et al., 2010; Ong Tone et al., 2021a). However, given the limited packaging capacity of scAAVs (Gruenert et al., 2016), we were unable to insert the Bcl-xL gene into scAAV2/5. To address this limitation, we inserted Bcl-xL into a single stranded AAV2/5 (ssAAV2/5) vector and investigated its transduction efficiency in FECD-SV-54F-73. We found that both ssAAV2/5-Bcl-xL and control ssAAV2/5 exhibited good transduction efficiencies in FECD-SV-54F-73 (54.54%±17.23 and 58.08%±12.30, respectively; *p*=0.788) (Figure 4A). We next investigated the ability of ssAAV2/5-Bcl-xL to protect FECD CECs from apoptosis by treating FECD-SV-54F cells with etoposide (Barcia et al., 2007; Fuchsluger et al., 2011a; Staehlke et al., 2024). We found that ssAAV2/5-mediated expression of Bcl-xL resulted in significant attenuation of etoposide-mediated apoptosis (23.96%±8.88) compared to ssAAV2/5 alone (70.43%±4.97; *p*=0.038) and non-transduced cells (71.68%±0.69; *p*=0.018) (Figure 4B, Supplemental Figure 4).

**Figure 4.**
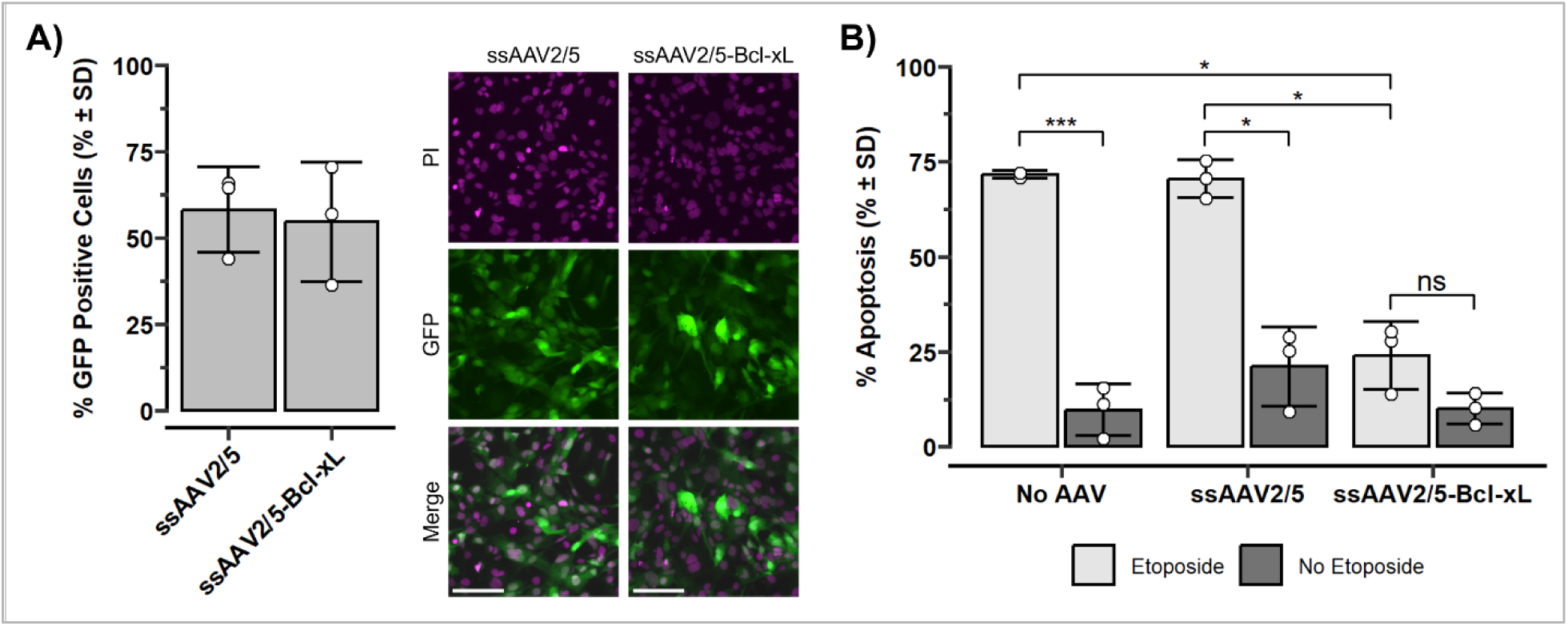
ssAAV2/5-Bcl-xL attenuates etoposide-mediated apoptosis in FECD. **(A**) Left: No significant differences were seen between ssAAV2/5 and ssAAV2/5-Bcl-xL transduction in FECD-SV-54F-73 (*p*=0.788). Transduction efficiency was quantified as the percentage of total cells expressing GFP. Statistical analysis was performed using Welch’s 2-sample t-test. Right: Representative images showing GFP expression in FECD-SV-54F-73 transduced with ssAAV2/5 or ssAAV2/5-Bcl-xL. Nuclei were stained using PI. Scale bar = 100 μm. **(B)** Quantification of apoptosis in FECD-SV-54F-73 CECs transduced with ssAAV2/5-Bcl-xL, ssAAV2/5-Bcl-xL, or no AAV, and treated with either etoposide or media only. Transduction with ssAAV2/5-Bcl-xL resulted in significantly lower levels of etoposide-induced apoptosis compared to control ssAAV2/5 (*p*=0.038) and non-transduced cells (*p*=0.018). Apoptosis was quantified as the percentage of total cells expressing Caspase-3/7. Statistical analysis was performed using 2-way ANOVA and post-hoc Tukey’s HSD. All experiments were done using N=3. AAV = adeno-associated virus, ss = single-stranded, GFP = green fluorescent protein, PI = propidium iodide. \**p* ≤ 0.05, \*\**p* ≤ 0.01, \*\*\**p* ≤ 0.001.

## 4. Discussion

In this study, we found that corneal endothelium from normal and FECD cadaveric donors expressed 12 different genes associated with AAV entry including three trans-membrane proteins, *KIAA0319L*, *GPR108*, and *TM9SF2*. KIAA0319L, also known as the Adeno Associated Virus Receptor (AAVR), has been identified as an essential cellular receptor that is required for transduction of several AAV serotypes, including the evolutionary distinct AAV2 and AAV5 (Meisen et al., 2020; Pillay et al., 2017). GPR108 is a highly conserved entry factor that plays a role in AAV entry for all major serotypes except AAV5 (Dudek et al., 2020; Meisen et al., 2020). Many AAV serotypes, including AAV2, require both AAVR and GPR108 to enter the cell. TM9SF2 is another critical factor for the transduction of multiple AAV serotypes, including AAV2, 5, 6, and DJ (Meisen et al., 2020). The remaining 9 genes identified in our bulk RNA sequencing (RNA-seq) dataset (*ACP2*, *ATP2C1*, *B2GAT3*, *C16orf62*, *EXT1*, *EXT2*, *NDST1*, *RGP1*, and *RNF121*) have also been implicated in AAV2 cellular binding, endosome/Golgi trafficking, endosomal escape, nuclear import, and/or gene expression (Rostami et al., 2023; Song et al., 2025). Taken together, this supports that AAV2 serotypes could be used for gene therapy targeting the CE.

Building on this RNA-seq data, the transduction efficiency and tropism of AAV2 can be further increased through alteration of the capsid. In addition to AAV binding genes deciding transduction, the capsid of an AAV is a major determinant of the vector’s tropism and transduction efficiency (Keswani et al., 2012; Vincent et al., 2014). Using pseudotype AAV2 vectors (where the genome of AAV2 is packaged inside the capsid of another) can lead to significantly greater transduction and altered tropism compared to either parental serotype (Keswani et al., 2012; Vincent et al., 2014). In AAV2/5 for example, the capsid of AAV2 is replaced with that of AAV5 to create AAV2/5, which contains the genome of AAV2 inside the capsid of AAV5. Many of these pseudotype vectors are well characterized in the literature and are often preferred over their parental serotypes for gene therapy applications (Burger et al., 2004).

AAV2, 5, 6, 8, and 9 have been explored in the CE of both human and non-human subjects (Bastola et al., 2020; Zhang et al., 2026). Zhang et al. demonstrated that intrastromal injections of AAV2, 6, 8, and 9 can transduce murine CECs in vivo, with AAV6 demonstrating the highest transduction efficiency. In rats, Lee et al. found that intracameral injections of AAV2, 5, and 8 demonstrated enhanced transduction in the CE, ciliary body, and iris. In rabbit corneas, Liu et al. found that AAV2/1, 2/2, 2/5, 2/7, and 2/8 can transduce all cell types of the cornea, including the epithelium, keratocytes, and endothelium. Using organ-cultured human corneas, Liu et al. also demonstrated that AAV2/1, 2/2, and 2/8 efficiently transduces all 3 cellular layers of the human cornea. Similarly, Hippert et al. showed that AAV2/1 and 2/8 can transduce ex vivo human corneas. These findings, combined with our bulk RNA-seq dataset, informed our selection of the 17 scAAV2 pseudotypes we tested in human CECs.

Following our bulk RNA sequencing, we screened 17 scAAV2 serotypes for transduction efficiency in human CECs and found that scAAV2/2 and scAAV2/5 resulted in the highest transduction efficiency in normal CEC and FECD cell lines, as well as in ex vivo specimens from healthy cadaveric donors and FECD patients undergoing endothelial keratoplasty.

Despite both scAAV2/2 and 2/5 demonstrating similarly high levels of transduction in CECs and tissues, there are several distinct differences in their structure and function. While both AAV2/2 and AAV2/5 share the same genome, that of AAV2, their capsids are quite different. AAV2 and AAV5 have evolutionarily distinct lineages and only share a 58% amino acid sequence identity (Pillay et al., 2017; Rayaprolu et al., 2013). During cellular entry and trafficking, AAV2 attachment to the host cell is largely mediated by heparan sulfate proteoglycans (HSPG), with several secondary receptors also playing a role (Pillay et al., 2017; Rayaprolu et al., 2013). Conversely, AAV5 utilizes sialic acid (SIA) moieties for their initial cellular attachment (Rayaprolu et al., 2013). The CE itself is abundant in both HSPG and SIA moieties, which likely influence the binding affinity of different AAV serotypes and favour both scAAV2/2 and scAAV2/5 despite their differences (Pillay et al., 2017; Zhang et al., 2026).

We have also demonstrated that ssAAV2/5-mediated expression of the anti-apoptotic molecule Bcl-xL significantly protected FECD human CECs from etoposide-induced apoptosis. Our findings are consistent with previous studies that have shown that lentiviral-mediated expression of Bcl-xL provides protection against apoptosis in mouse CECs and corneal grafts in vitro (Barcia et al., 2007). Similarly, lentiviral-mediated expression of Bcl-xL has been explored as a strategy to promote human cadaveric donor corneal tissue survival during storage, as well as to attenuate induced apoptosis in cultured human CECs and ex vivo tissues (Fuchsluger et al., 2011b).

Bcl-xL has been shown to be a neuroprotective factor in several neurodegenerative conditions, such as amyotrophic lateral sclerosis (ALS), Alzheimer’s, and Parkinson’s diseases (Garrity-Moses et al., 2005). Garrity-Moses et al. demonstrated that AAV-mediated overexpression of Bcl-xL significantly reduced motor neuron death in rat ALS in vitro models through inhibition of excitotoxicity-induced apoptosis. Matsuoka et al. found that adenovirus-mediated expression of Bcl-xL in vitro enhanced the survival of cortical neurons in a primary neuronal rat model of Alzheimer’s disease. Similarly, Kaspar et al. showed that retrograde AAV2 in vivo delivery of Bcl-xL protected mature entorhinal projection neurons from induced cell death in rat studies of Alzheimer’s and Parkinson’s diseases. While there is debate about whether rescuing cells by interrupting the apoptotic cascade is sufficient to relieve the symptoms associated with degenerative diseases, there is strong evidence suggesting that preserving mitochondrial function is a pertinent approach to the long-term inhibition of neurodegeneration (Malik et al., 2005). Mas-Bargues et al. demonstrated that Bcl-xL overexpression preserves mitochondrial structure and function in mouse T cells, thereby aiding in the regulation of intrinsic apoptosis and oxidative stress, increasing resistance to stress-induced apoptosis. The protective effect that Bcl-xL overexpression has in both neurodegenerative disease and mitochondrial function, combined with its significant role in the apoptotic cascade, make Bcl-xL overexpression a promising strategy when considering the modulation of cell fate in neurodegenerative diseases and FECD (Malik et al., 2005; Mas-Bargues et al., 2025; Zhu et al., 2014).

An important limitation of Bcl-xL as a therapeutic is that it does not protect against non-apoptotic cell death. While oxidative stress-induced apoptosis has been identified as a significant cause of cell death in FECD, other non-apoptotic mechanisms leading to cell death such as iron-mediated lipid peroxidation, ferroptosis, necrosis, and parthanatos have been reported (Saha et al., 2024; Sakakura et al., 2023). These cell death mechanisms are not dependent on Bcl-xL, thereby limiting the effectiveness of Bcl-xL-based gene therapy approaches at preventing this type of cell death (Bleicken et al., 2017; Dou et al., 2021; Saha et al., 2024; Sakakura et al., 2023; Westphal et al., 2014).

Additionally, while our ssAAV2/5-Bcl-xL construct shows potential as a therapeutic strategy in the prevention of apoptosis associated with FECD, this approach may prolong the life of diseased FECD CECs and may not rescue dysfunctional CECs. Additional studies are needed to further explore FECD pathogenesis and how targeting the anti-apoptotic pathway can be developed as part of a therapeutic strategy. One possibility may be using ssAAV2/5-Bcl-xL to attenuate apoptosis before rescuing these diseased FECD cells with gene therapy and/or an antioxidant compound to restore functionality. While the clinical translational potential of our ssAAV2/5-Bcl-xL construct is promising, further explorations are required to investigate ssAAV2/5-Bcl-xL tropism in other ocular tissues prior to clinical application.

We also did not observe a significant increase in apoptosis associated with AAV transduction in CECs. This is an important finding as potential gene therapies should not induce apoptosis in CECs. Our findings support that AAVs are a viable method for the ex-vivo delivery of anti-apoptotic genes and/or gene therapy to the CE, but requires further exploration to optimize in vivo delivery.

In vivo, human CECs are arrested in the G1 phase of the cell cycle and possess little regenerative or proliferative capacity (Benischke et al., 2017). Immortalized human CECs, however, are proliferative, and continuously divide. While AAVs can infect both dividing and non-dividing cells, the long-term effects differ between the cell types (Daya and Berns, 2008; Schultz and Chamberlain, 2008). Since AAVs are non-replicating and do not integrate into the host genome, as cell division occurs in cultured CECs, the viral DNA is lost, and transduction ceases (Daya & Berns, 2008). As the host cell divides, the AAV genome is not replicated alongside the cellular DNA, leading to a dilution of the vector genome in the subsequent daughter cells (Schultz and Chamberlain, 2008). This dilution of the AAV vectors also leads to decreased long-term gene expression in dividing cells. Therefore, in cultured CECs, the transgene expression of AAVs is transient and gradually decreases as the cells divide (Schultz and Chamberlain, 2008). This dilution and decreased long-term transgene expression is not a major concern in the corneal endothelium, where there is significantly decreased proliferative capacity in vivo (Schultz and Chamberlain, 2008).

AAVs can be packaged as either single-stranded (ss) or self-complementary (sc) constructs. In ssAAVs, the genome is packaged as a linear ssDNA molecule approximately 4.7kb in length (Gruenert et al., 2016). The single-stranded nature of the DNA requires the host cell to convert the ssDNA into a double-stranded (ds) DNA template prior to expression. This synthesis is a rate-limiting step for ssAAV transduction (Gruenert et al., 2016). scAAVs bypass this synthesis step, as these vectors contain dimeric inverted repeat regions that can spontaneously fold into dsDNA templates, leading to a faster onset of gene expression and higher transduction efficiencies (Gruenert et al., 2016; Mohan et al., 2005; Parker et al., 2009; Sharma et al., 2010). The self-complementary sequence, however, limits the packaging capacity of scAAVs to 2.3kb, approximately half that of ssAAVs (Gruenert et al., 2016). In the initial screening and selection of AAV2 serotypes, scAAV2 serotypes were utilized. Due to the limited packaging capacity of scAAVs, however, the anti-apoptotic construct containing Bcl-xL was unable to fit in the scAAV2 construct utilized in the initial serotype screening (Mohan et al., 2005). To accommodate the packaging size requirements of Bcl-xL, a ssAAV2/5 construct was used. The anti-apoptotic ssAAV2/5-Bcl-xL and control ssAAV2/5 constructs were tested in FECD-SV-54F-73 human CECs and we found that viral transduction was comparable to scAAV2/5. We then demonstrated the ability of ssAAV2/5-mediated expression of Bcl-xL to protect against etoposide-induced apoptosis in FECD CECs.

## 5. Conclusions

Our study demonstrates the ability of various AAV2s to transduce the corneal endothelium, both from healthy cadaveric donors and FECD patients, and provides a framework in which other AAV-based gene therapies can be designed for the corneal endothelium. The significant attenuation of apoptosis seen in FECD CECs transduced with ssAAV2/5-Bcl-xL shows promise as a potential therapeutic in the treatment of FECD. Furthermore, ssAAV2/5 can be used to deliver targeted gene therapies to the human CE, and future studies may utilize this framework to deliver alternative gene therapies to the corneal endothelium. In addition to FECD therapeutics, ssAAV2/5-Bcl-xL has potential applications in extending the viability of healthy cadaveric donor corneas in eye banks. By protecting CECs against that apoptosis associated with extended storage, there is potential to extend the donor tissue transplantation window, therefore increasing the number of donor corneas available for corneal transplantation. However, future studies are needed to explore this possibility.

## CRediT authorship contribution statement

**Ness Little:** Data Curation, Formal Analysis, Investigation, Methodology, Project Administration, Software, Validation, Visualization, Writing – Original Draft, Writing – Review & Editing.

**Judy Yan:** Investigation, Methodology, Project Administration, Resources, Supervision, Validation, Writing – Review & Editing.

**Narisa Dhupar:** Investigation.

**Stephan Ong Tone:** Conceptualization, Funding acquisition, Methodology, Project Administration, Resources, Supervision, Validation, Writing – Review & Editing.

## Funding

This work was supported by Fighting Blindness Canada.

## Declaration of competing interest

This work was supported through a Fighting Blindness Canada Research Grant. Dr. Ong Tone has the following disclosures: Labtician (research grant/financial support), Rx Renewal (consultant/consulting fees), and Sun Pharma (consultant/consulting fees).

## Acknowledgements

We would like to thank Fighting Blindness Canada, The Sunnybrook Foundation, and The Vision Science Research Program for funding the study. We would like to thank Dr. Ula Jurkunas at the Schepens Eye Research Institute in Boston, Massachusetts for graciously providing the cell lines.

## List of Abbreviations

AAV: Adeno-associated virus
AAVR: Adeno-associated virus receptor
ALS: Amyotrophic lateral sclerosis
Bcl-xL: B-cell lymphoma-extra large
CE: Corneal endothelium
CEC: Corneal endothelial cell
DM: Descemet’s membrane
DMEK: Descemet membrane endothelial keratoplasty
ds: Double-stranded
ECM: Extracellular matrix
FECD: Fuchs Endothelial Corneal Dystrophy
FPKM: Fragments per kilobase of transcript per million fragments mapped
GFP: Green fluorescent protein
HSPG: Heparan sulfate proteoglycan
MOMP: Mitochondrial outer membrane permeabilization
NGS: Next generation sequencing
PBS: Phosphate-buffered saline
PI: Propidium iodide
RIN: RNA integrity number
RNA-seq: RNA sequencing
sc: Self-complementary
SIA: sialic acid
ss: Single-stranded

## Data Statement

Data will be made available on request.

