## Supplemental Figures for "Adeno-Associated Virus Mediated Expression of Bcl-xL Attenuates Apoptosis in Fuchs Endothelial Corneal Dystrophy"

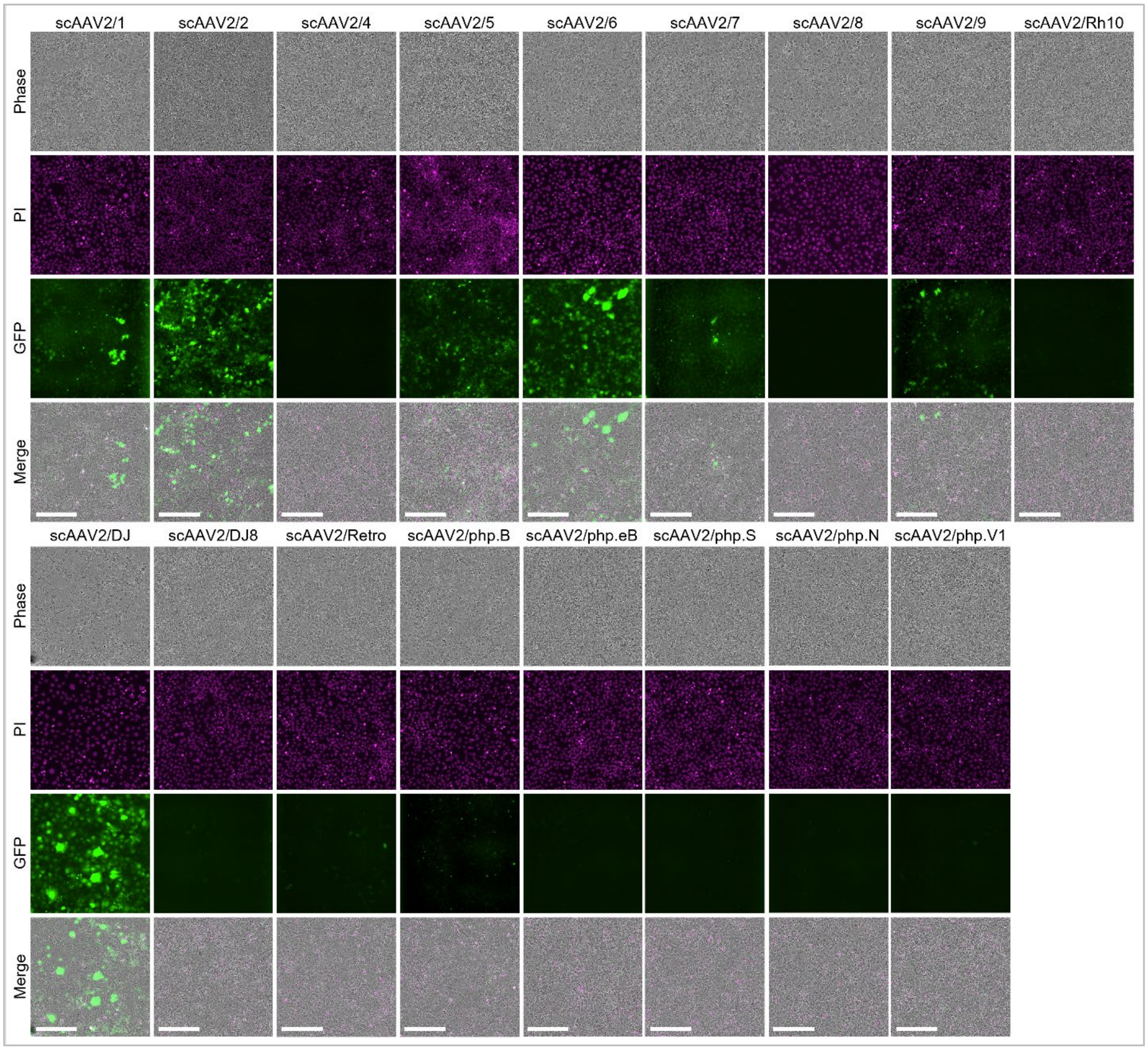


**Supplementary Figure 1.** Screen of 17 scAAV2 serotypes in HCEC-21T cells for GFP expression. scAAV2/2, 2/5, 2/6, 2/DJ demonstrated the highest transduction efficiencies of the seventeen serotypes (scAAV2/1 (6.46%±3.54), scAAV2/2 (63.81%±5.66), scAAV2/4 (0.09%±0.06), scAAV2/5 (37.95%±14.17), scAAV2/6 (33.61%±14.13), scAAV2/7 (2.74%±0.76), scAAV2/8 (12.01%±7.90), scAAV2/9 (8.24%±0.58), scAAV2/Rh10 (3.47%±1.27), scAAV2/DJ (37.64%±11.78), scAAV2/DJ8 (7.51% ±3.89), scAAV2/Retro (7.28%±1.58), scAAV2/php.B (2.13%±1.82), scAAV2/php.eB (1.70%±0.32), scAAV2/php.S (2.08%±1.43), scAAV2/php.N (0.60%±0.10), scAAV2/php.V1 (1.10%±0.26)). Transduction efficiency was quantified as the percentage of total cells expressing GFP. All experiments were done using N=3, except scAAV2/php.N and scAAV2/php.V1 (N=2). AAV = adeno-associated virus, sc = self-complimentary, GFP = green fluorescent protein, PI = propidium iodide. Scale bar = 200 μm.

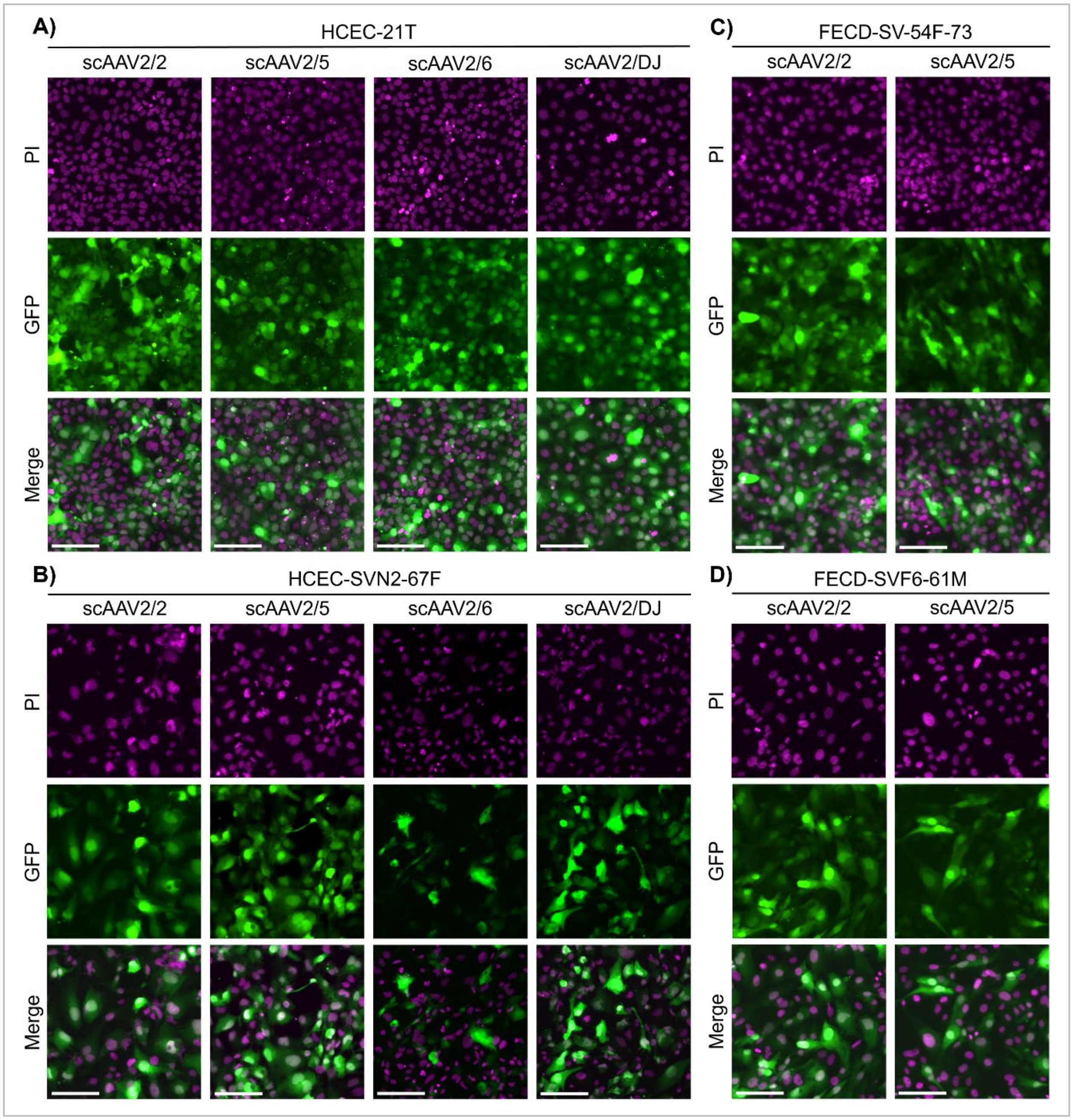


**Supplementary Figure 2.** scAAV2 serotypes can transduce normal and FECD HCECs with high efficiency. scAAV2 transduction in **(A)** HCEC-21T, **(B)** HCEC-SVN2-67F, **(C)** FECD-SV-54F-73, **(D)** FECD-SVF6-61M cells. Nuclei were stained using PI. All experiments were done using N=3. AAV = adeno-associated virus, sc = self-complimentary, GFP = green fluorescent protein, PI = propidium iodide. Scale bar = 100 μm.

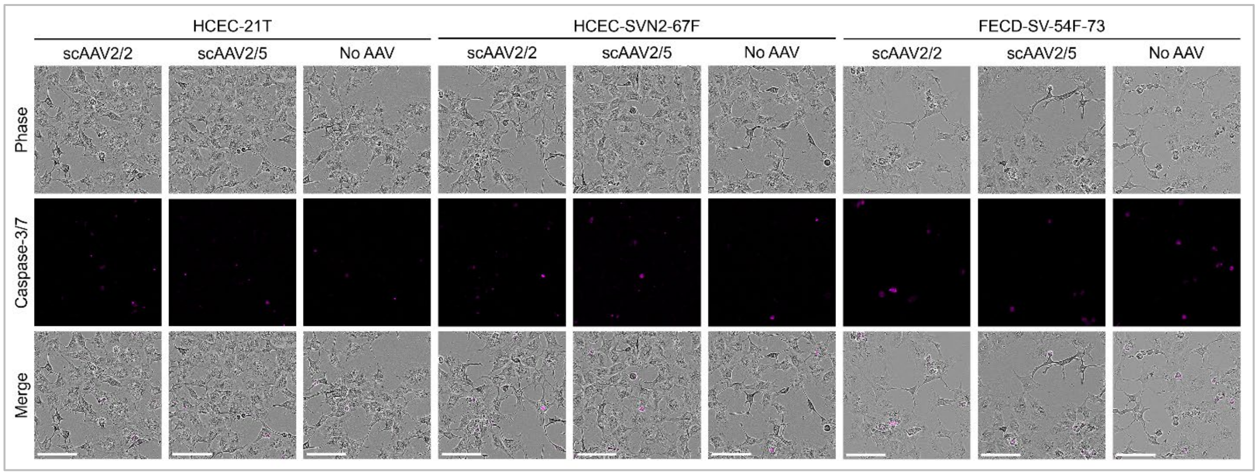


**Supplementary Figure 3.** scAAV2 transduction does not increase apoptosis in CECs. Representative live cell images showing levels of apoptosis in **(A)** HCEC-21T, **(B)** HCEC-SVN2-67F, and **(C)** FECD-SV-54F-73 cells transduced with scAAV2/2 and scAAV2/5. Total cells were visualized via phase; apoptosis was visualized via Caspase-3/7 staining. All experiments were done using N=3. AAV = adeno-associated virus, sc = self-complimentary. Scale bar = 100 μm.

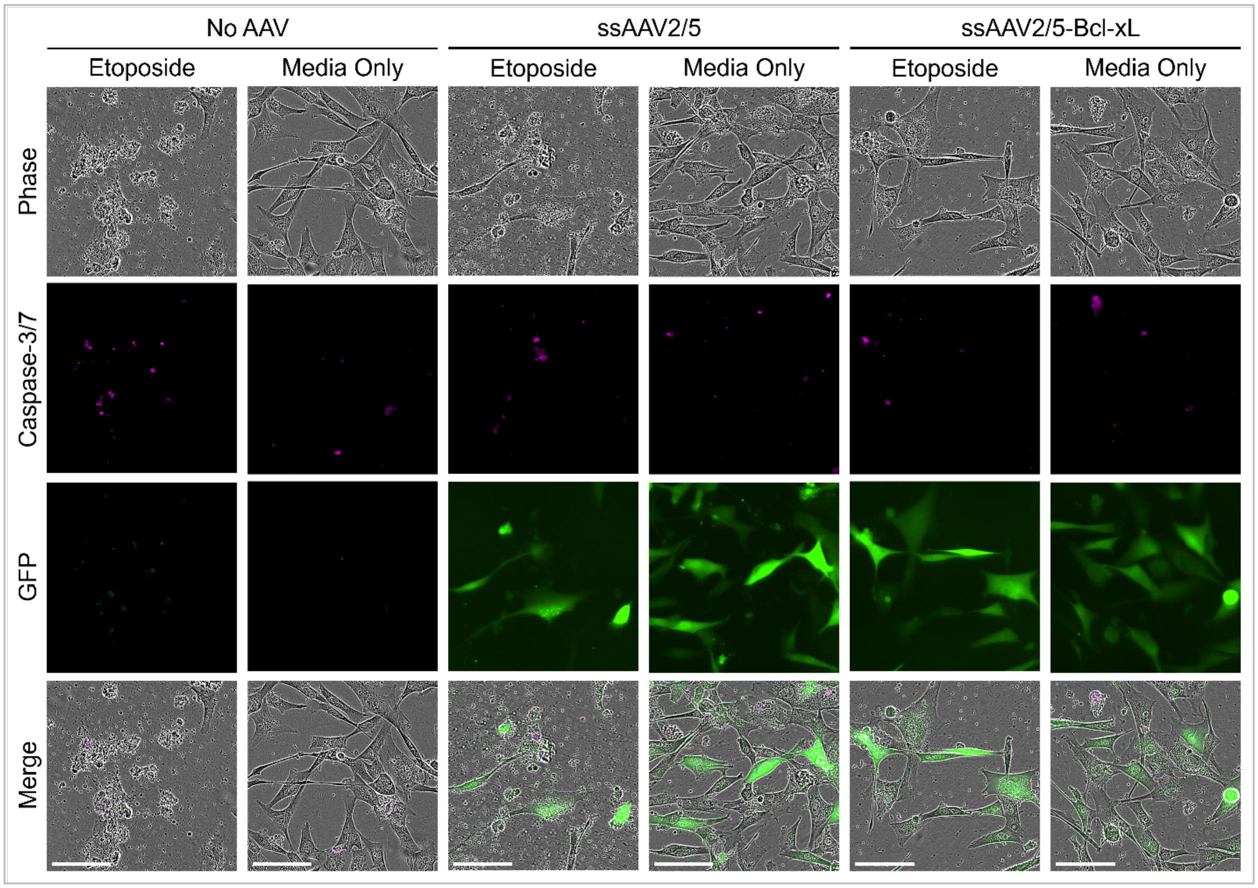


**Supplementary Figure 4.** ssAAV2/5-Bcl-xL attenuates etoposide-induced apoptosis in FECD. Representative live cell images showing the levels of apoptosis in FECD-SV-54F-73 cells transduced with no AAV, ssAAV2/5, or ssAAV2/5-Bcl-xL. Total cells were visualized via phase. Apoptosis was visualized via Caspase-3/7 staining. GFP was used to validate AAV transduction. All experiments were done using N=3. AAV = adeno-associated virus, ss = single-stranded. GFP = green fluorescent protein. Scale bar = 100 μm.
